# Projection Statistics – ProST: Online statistical assessment of group separation in data projection analysis

**DOI:** 10.1101/2024.09.04.611273

**Authors:** Danny Salem, Anuradha Surendra, Graeme SV McDowell, Miroslava Čuperlović-Culf

## Abstract

**Motivation:** Unsupervised data projection for the determination of trends in the data, visualization of multidimensional data in a reduced dimension space or feature space reduction through combination of data is a major step in data mining. Methods such as Principal Component Analysis or t-Distribution Stochastic Neighbor Embedding are regularly used as one of the first steps in computational biology or omics investigation. However, the significance of the separation of sample groups by these methods generally relies on visual assessment. User-friendly application for different projection methods, each focusing on distinct data properties, are needed as well as a rigorous method for statistical determination of the significance of separation of groups of interest in each dataset.

**Results:** We present Projection STatistics (ProST), a user-friendly solution for data projection analysis providing three unsupervised (PCA, t-SNE and UMAP) and one supervised (LDA) approach. For each method we are including a novel statistical investigation of the significance of group separation with Mann-Whitney U-rank or t-test analysis as well as necessary preprocessing steps. ProST provides an unbiased, objective application of the determination of the significance of the separation of measurement groups through either linear or manifold projection analysis with methods ranging from a focus on the separation of points based on major variances or on point proximities based on distance.

**Availability:** The ProST software application is freely available at https://complimet.ca/shiny/ProST/ with source code provided on https://github.com/complimet/prost.

**Supplementary information:** Supplementary help pages are provided at https://complimet.ca/shiny/ProST/.

## 1 Introduction

Dimension reduction methods are one of the main tools in bioinformatics used for presenting information about trends in data, often through insightful visualizations. In fact, an overview of data using projection methods is often seen as a first step in analysis as it provides unbiased, unsupervised, fully data-driven information. Principal Component Analysis (PCA) is arguably the most utilized method in this group providing information about the variation in the data represented by a projection on orthogonal axes through a linear combination of variables (Dorrity et al. 2020). T-Distribution Stochastic Neighbor Embedding (t-SNE) is an increasingly popular method for dimensionality reduction. It is a projection of a datapoints in a high-dimensional space onto a low-dimensional space that aims to maintain the proximity of nearby points (van der Maaten et al. 2008). Uniform Manifold Approximation and Projection (UMAP) method aims to devise a low dimensional representation of the multidimensional data on estimated topology that preserves local relationships in the data (McInnes, and L, Healy, 2018). Linear discriminant analysis (LDA) is a supervised technique that performs dimensionality reduction by fitting linear combinations of the variables in a dataset that optimally group datapoints within classes together and separate datapoints from different classes apart. LDA is based on Fisher’s linear discriminant developed in the 1930s by Sir Fisher. All these approaches provide a unique perspective on datasets with a focus either on the determination of major variances in the data and searching for the projection leading to largest sample separation (PCA), exploring similarities between points (t-SNE, UMAP) or distances (LDA). The final assessment of sample group separation is generally based on visual inspection of 2D or 3D plots provided by these methods. Although PCA is widely available through some excellent user-friendly tools (Pang, et al. 2024), seamless, user-friendly access to other projection methods is lacking. More importantly, with projection methods used to provide information about the data-driven, unsupervised separation of sample or feature groups, there is a need for a statistical measure of the level of group separation resulting from the dimension reduction methods. Statistical information can provide an unbiased measure of the significance of observed separation of groups of interest, thus enhancing understanding of the data properties explored with different projection methods.

To address these needs, we are providing a new open-access web-based application called PROjection STatistics – ProST, that includes a user-friendly application of PCA, UMAP, t-SNE – the unsupervised projection methods as well as LDA – supervised alternative, with statistical information about the statistical significance of sample groups and cohorts, as well as readily available visualization of results.

## 2 Implementation

ProST is implemented in Python with a RShiny front-end. It is compatible with all web browsers. Users can choose between four different dimensionality reduction methods including: PCA, t-SNE, UMAP and

LDA.

Briefly:

1. PCA is performed using the *sklearn* Python library. PCA uses Singular Value Decomposition (SVD) of the data to determine lower dimensional projections. The main variances in the data are represented in Principal Components 1 and 2 and they are shown in ProST.
2. T-SNE is implemented using the openTSNE Python library (PoliČar et al. 2019). T-SNE aims to provide an embedding of high-dimensional data into low dimensions by preserving the proximity between neighbouring points and the distance between distant points. Point proximity is assessed through the analysis of Gaussian distribution of point distances with the distribution’s standard deviation dependent on the perplexity hyperparameter. In ProST, the user is able to take advantage of all the recent improvements and best practices that have been implemented to improve the t-SNE algorithm (Kobak and Berens 2019). These include PCA initialization, optimal learning rate (Belkina et al. 2019), multiscale perplexity and different affinity metrics. The main aim of these techniques is to substantially improve global structure reconstruction while maintaining the quality of the local structure. It also uses an accelerated gradient calculation algorithm called FIt-SNE (Linderman 2019) to improve the execution time of t-SNE optimization. t-SNE in ProST allows users to either use preselected hyperparameters or to select between Euclidean or Cosine distance, and different values for parameters such as perplexity, E.E. coefficient and iterations as well as number of iterations. Detailed consideration of the effect of these parameters on t-SNE result is presented previously (Belkina et al. 2019, Kobak and Berens 2019).
3. UMAP is focused on maintaining local groupings of similar points while preserving global structures between more distant observations (Dorrity, et al. 2020). UMAP makes the assumptions that points in a dataset are uniformly distributed on a Riemannian manifold, that the Riemannian metric can be approximated as locally constant and that the manifold is locally connected. In this case, the embedding is determined by searching for a low dimensional projection that has the closest fuzzy topological structure. Global structure information is arguably better preserved in UMAP projections (McInnes and Healy 2018) than in t-SNE. UMAP is implemented using the Python library *umap* (McInnes et al. 2018). In ProST users can define the number of neighbours and minimum distance prior to UMAP analysis. The minimum distance refers to the smallest possible distance between points in mapping produced by UMAP. Changing the minimum distance will affect how closely points will be packed in the 2D representation. Selection of the number of neighbours defines the size of the neighbourhood around each point used to estimate the manifold. The number of neighbours parameter represents a trade-off between local and global structure. The opportunity to select the number of neighbours and the minimum distance aims to help users to ensure that UMAP preserves local and global structures.
4. LDA is a supervised dimensionality reduction method that, similarly to PCA, projects the data onto new axes, however, the LDA projection focuses on maximizing variability between the classes and reducing the variability within the classes. As a linear and supervised projection method LDA finds a straight line or a plane that optimally separates groups in the input set. LDA assumes a normal data distribution and identical covariance matrices between classes. The main consideration in LDA is that the dimensionality of the projected set has less than number of sample groups -1. Thus, for LDA to provide the projection the dataset has to have at least 3 distinct group, i.e. class labels.

Prior to the projection analysis, user can optionally select to impute data using one of several standard approaches (1/5 minimal value, median value, KNN (Hastie, et al. 1999; Troyanskaya, et al. 2001) or MICE (van Buuren, et al. 1999; van Buuren et al. 2006)) as well as normalize and/or log transform data.

Statistical analysis is performed for the selected projection method using either a Mann-Whitney U rank test (using option *mannu* in Python library *scipy*) or a t-test. The Mann-Whitney U rank test is a preferred option as projection values cannot be assumed to have normal distribution. In this case we are determining the statistical significance that the distribution of projection values in two coordinates for different groups of samples is different. Analysis is done pairwise, with a p-value provided for comparison of each pair of groups. The value for each sample in this analysis is its projection measure in either dimension 1 or 2 as provided by the presented projection method. Statistical analysis compares the significance of the differences between projections in dimensions 1 and 2 between each pair of sample groups. The distribution of values is shown as a violin plot and p-values for each pairwise comparison are shown in the table. Provided tables show data from each step in the analysis from information about the input data (“Input Data”), data following all selected data transformations including imputation, normalization and/or log transformation (in “Transformation Data” table); projection values for the first two components (in “Reduction Table”) and finally p-values for pairwise group comparisons for two projections (in “Statistics Table”). All tables and plots can be downloaded.

A detailed user manual with software information is provided on the web site: https://complimet.ca/shiny/ProST/.

## 3 Results

The ProST application settings are conceptually divided into three blocks (Figure 1A):

**Figure 1.**
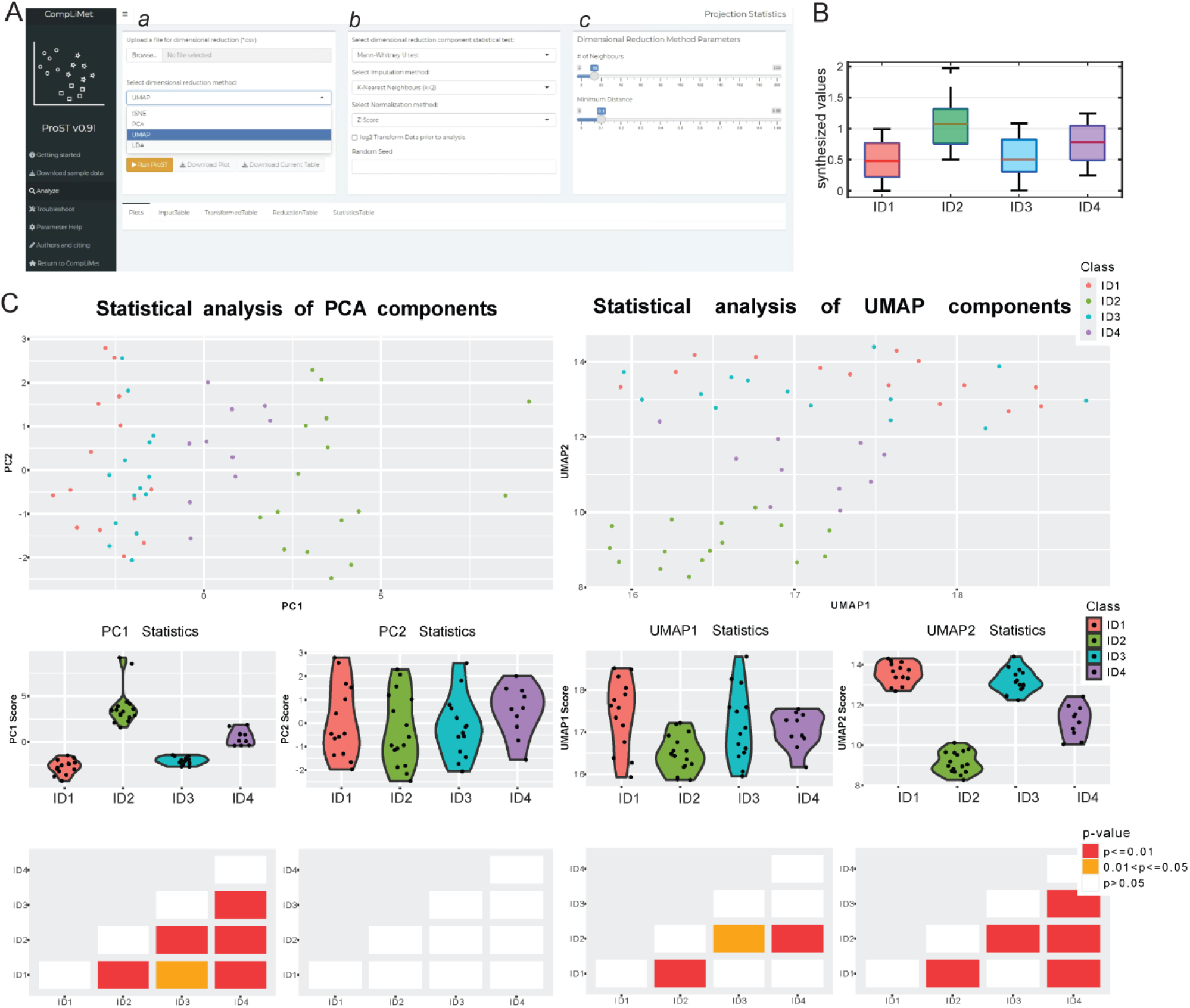
A. Analysis window of ProST showing options for the projection methodologies, data preprocessing and method hyperparameters. Available at https://complimet.ca/shiny/ProST/; B. Simulated dataset includes 60 samples with four cohorts and 20 features. Each simulated cell includes random values between 0 and 1. ID1 values are only random values between 0 and 1; ID2 values have 0.5 added to each cell; ID3 adds 0.1 to 14 features in ID1, and new random values for the other 6 features; ID4 adds 0.25 to random values. Shown is a boxplot of values for all features across simulated samples for four groups. C. PCA and UMAP results for z-score normalized values in the simulated set. Shown are results produced by ProST example projection plots and the level of resulting separation. Mann-Whitney U-test calculated p-values between pairs of sample groups are shown schematically. P<0.01, indicating major group separation, is obtained for the separation between all sample groups except for groups ID1 and ID3, where p-value is between 0.01 and 0.05 as is expected due to smaller difference between these two groups.

a. *Input information upload block* – the dataset file and associated information is entered by the user in this block. The input file has to be in .CSV format with samples in rows and features in columns. The user specified column with sample class information will be used for statistical analysis of the projection results as well as sample labelling. This column can be anywhere in the input file and user can select column from the drop-down list of all columns in the input file. Analysis will be done using all numeric data columns and thus other text information columns, e.g. sample labels or other sample information, can be included and will not affect the analysis. A minimum of two features are required. Alternatively, users can select to not use labels for the dimensional reduction analysis in which case projection will be shown but there will not be sample class analysis. Mann-Whitney test does not have sample size requirements so any number of samples in a cohort can be analyzed using this test. In this step users can choose between four provided projection methods.
b. *Statistical analysis of components and data transformation block* – statistical significance of the projection sample group separation can be established using either Mann-Whitney U test or two independent sample t-test and this selection is made in this block. Additionally, there are options for users to impute missing data from the dataset using one of four provided methods and to normalize data using one of five provided methods. Data can additionally be log transformed if desired.
c. *Dimension reduction method parameters block* – hyperparameters for t-SNE and UMAP projection methods can be selected here by the user. For t-SNE the user is provided with an option to change the distance metric, the perplexity, the affinity metric, the number of optimization steps and the early exaggeration (EE) coefficient. For UMAP the user may select the number of neighbours and minimum distance. The default values for these parameters will work for many use cases, however they can have a fundamental impact on the projection result and ideally users should try several different values using the parameter help section of the website as a guide.

The utility of ProST is shown using a simulated toy example (Figure 1B and C). The dataset used in the example includes 20 features with random values in four groups of samples. Group 1 (ID1) includes random values between 0 and 1 for all samples and features; Group 2 (ID2) adds 0.5 to random values between 0 and 1 for all cells, Group 3 (ID3) is simulated by adding 0.1 to values in ID1 for 14 features and for 6 features has new random values for all samples; Group 4 (ID4) adds 0.25 to random values from 0 to 1. This simple simulation shows four groups of samples with significant differences between groups 1 and 2 and very limited differences between values in groups 1 and 3. Figure 1B shows overall data spread in these three simulated groups. Figure 1C shows examples of analysis of this dataset using methods provided in ProST with statistical representation of the significance of group separation in the projection space. Data is auto-scaled (z-score normalized). Major sample group separation is clearly visible in the projection plots with p-value and violin plot of projection values for each sample group additionally indicating statistical significance of the separation. Shown are results obtained using UMAP and PCA as examples. Analysis shows statistically significant difference, p<0.01, in the separation of all groups in PC1 except between ID1 and ID3, with p<0.05. In PCA all variances between samples are in PC1 with no statistically significant separation in PC2. Furthermore, the UMAP projection analysis shows high level of significance of the separation between ID2 and ID1 and ID4 at p-value < 0.01 in the first projection, and similarly to PC1, p<0.01 for all groups except group ID1 vs. ID3 in the second UMAP dimension. In this analysis ProST provides the projections as well as further information about the statistical significance with tabular information in the Statistics Table tab showing exact p-values for the statistical significance of projections in two dimensions between all sample groups. Difference in the separation between groups is clearly visible and calculated p-values agrees with the simulated level of difference between sample groups.

## 4 Conclusion

ProST is an open-access Web-based application for the data visualization and statistical representation of projection sample separation. Primary use is for omics data analysis although this is a general application that can be used in principle for any numeric data. ProST and detailed instruction for use are available at https://complimet.ca/shiny/ProST/. Future development of ProST will include the addition of more projection methods as well as further development of graphical representation of results.

## Funding and Acknowledgements

This work was supported in part by operating grant AI-4D-102-3 from the NRC AI4D Program. Authors would like to thank Dr. Steffany Bennett from University of Ottawa for her collaboration on the https://compliment.ca site. *Conflict of Interest:* none declared.

## Notes

https://complimet.ca/shiny/ProST/

### Competing Interest Statement

The authors have declared no competing interest.

